# Inter-modality source coupling: a fully-automated whole-brain data-driven structure-function fingerprint shows replicable links to reading in a large-scale (N∼8K) analysis

**DOI:** 10.1101/2024.03.07.583896

**Authors:** Aline Kotoski, Jingyu Liu, Robin Morris, Vince Calhoun

## Abstract

**Objective:** Both structural and functional brain changes have been individually associated with developing cognitive processes such as reading. However, there is limited research about the combined influence of resting-state functional and structural magnetic resonance imaging (rs-fMRI and sMRI) features in reading development, which could provide insights into the interplay between brain structure and function in shaping cognitive growth. We propose a method called inter-modality source coupling (IMSC) to study the coupling between the rs-fMRI and sMRI and its relationship to reading ability in school-age children.

**Methods:** This approach is applied to baseline data from four thousand participants (9-11 years) and replicated in a second group. Our analysis focused on the relationship of IMSC to overall reading score.

**Results:** Our findings indicate that higher reading ability was linked with increased function-structure coupling among higher-level cortical regions, particularly those links between the inferior parietal lobule and inferior frontal areas, and conversely, lower reading ability was associated with enhanced function-structure coupling among the fusiform and lingual gyrus. Our study found evidence of spatial correspondence between the data indicating an interplay between brain structure and function in our participants.

**Conclusion:** Our approach revealed a linked pattern of whole brain structure to the corresponding functional connectivity pattern that correlated with reading ability. This novel IMSC analysis method provides a new approach to study the multimodal relationship between brain function and structure.

**Significance:** These findings have interesting implications for understanding the multimodal complexity underlying the development of the neural basis for reading ability in school-aged children.

## I. Introduction

Reading requires the organization and coordination of multiple areas and functions in the brain as a developing, complex higher-order cognitive skill [1]. Studying reading development in children is a fundamental aspect of education research and practice. Its significance is underlined by recognizing that the ability to read proficiently is not only a crucial life skill but also provides a gateway to acquiring knowledge across various subjects [2]. Importantly, many studies have found that children with early difficulties in learning to read during initial grades often have persistent reading challenges as they get older [3]. These results emphasize the importance of studying reading development in children as it significantly influences their educational trajectory. Unfortunately, the precise neural processes and mechanisms underlying reading development are not fully understood [4]. Multiple ongoing research efforts continue to explore how different brain areas and functions contribute to various aspects of reading and its components, including decoding, comprehension, and fluency [4, 5].

Over the past two decades, magnetic resonance imaging (MRI) has been widely used as a non-invasive method to study the brain’s spatial and temporal attributes [6]. Functional MRI (fMRI) has been used to indirectly measure the brain’s condition and activity by measuring the changes in blood-oxygen levels [6]. The rs-fMRI applies this technique at rest and investigates the connectivity of brain regions where no task or stimulus is being applied to the participant [7]. sMRI has provided information about different tissue attributes and structure within the brain, such as white and gray matter integrity [8]. Both rs-fMRI and sMRI have also been extensively used to study reading development in children [8-12]. As an example, several studies have shown that there is an increase in whole brain volume and white matter coherence during its development [8, 13-15], and such cortical development has been related to reading development [8].

A major goal of modern neuroscience is understanding the linkages between such brain development and cognitive development [16]. There have been many structural and functional brain attributes that have been individually linked to a wide variety of basic and higher order cognitive processes, including reading. As an example, a recent study utilized inter-subject correlations and identified a widespread set of brain regions, including the anterior cingulate and precuneus, that differentiated groups of good or poor readers [17]. However, despite these many individual reported associations between neural structures using sMRI and related methods, or neural functions using rs-fMRI approaches, and cognition, there has been little work evaluating the linked or associated relationships between rs-fMRI and sMRI and such linkages’ relationship to the development of reading. Studying these critical brain attributes in combination provides the foundation on which to examine the complexities of how brain function and structure interact during development, and how this intersection may jointly influence reading development.

Here we propose a novel method called inter-modality source coupling (IMSC) to study the relationship between the function (using rs-fMRI) and structure (using sMRI) of the brain in children and how this integrating analysis relates to a measure of reading ability. To do this we proposed a novel approach that involved a spatially constrained independent component analysis (ICA) applied to both rs-fMRI and sMRI data which allowed us to identify linked brain networks adapted to the individual modality whereas also offering a semi-blind, data-driven index of correspondence across modalities of these brain networks. We then leverage this correspondence to estimate coupling between the connectivity matrix from rs-fMRI data and the loading parameter vectors from sMRI data. We next use the IMSC approach to study structural/functional links to reading ability. Our analysis revealed significant differences in whole-brain function, structure, and the relationship with reading ability, which were replicated in a second sample. Notably, we observed a correlation between reading ability and multiple brain areas, all positive correlations concentrated in frontal brain areas, such as the superior and inferior parietal lobules, whereas negative correlations were centered in early processing areas, such as the fusiform and lingual gyrus.

## II. Methods

The Adolescent Brain and Cognitive Development (ABCD) study is a 10-year longitudinal study that evaluates the development of children [13]. This study was undertaken, in part, to examine the linkage between brain structure and function, especially during the developmental period, which is complex and understudied. One of the challenges in such an undertaking has been to identify a common framework for linking the two. Most studies to date have focused on fixed brain regions of interest, however such approaches make rather strong assumptions that the pre-defined regions represent homogeneous aspects of the brain data and/or the functions of interest. Here we propose a novel multimodal fingerprint analysis method called IMSC, in an attempt to provide a more data-driven correspondence between functional and structural MRI (sMRI and rs-fMRI) using participants from the ABCD study.

We then utilize our approach to study the relationship of our structure/function coupling measures to a measure of reading ability by analyzing data from four thousand participants (9-11 years) from the ABCD dataset. We also replicate our results in another set of four thousand participants. Note, here we define the (computational) reproducibility as the evaluation of the results using the same data and using the same code, and the replicability as evaluation of the results on different data using the same code [18]. For the reading ability analysis, we used a score from the NIH Toolbox Oral Reading Recognition Test Age 3+ v2.0, the Theta Individual Ability Score (which is not age standardized and represents a child’s relative reading ability on the measure). This test is used to measure reading decoding skills and evaluate individual reading ability, asking the participants to pronounce single words [19, 20]. The sample, overall design, and detail on ethical considerations of the ABCD study are described by [21-23].

### A. Data

This study included eight thousand participants, that were separated into two different analyses (first analysis/replication analysis). The first analysis group with four thousand participants (1926 females) were aged between 8 years and 11 months and 11 years and 0 months (mean = 9 years and 11 months; SD = 7 months). The replication group included another set of four thousand participants (1916 females) also aged between 8 years and 11 months and 11 years and 0 months (mean = 9 years and 11 months; SD = 7 months).

### B. MRI parameters

Data were collected on three types of 3T scanners (Siemens Prisma, General Electric (GE) 750 and Philips) all with a standard adult-size head coil. The MRI sequences acquired and used for this work were a T1 structural scan (TR = 2500/2500/6.31 ms respectively; TE = 2.88/2/2.9 ms respectively); and a resting state sequence. The resting-state EPI sequence used the following parameters: TR = 800 ms, TE = 30 ms, 60 slices, flip angle = 52o, matrix size = 90 x 90, FOV = 216 x 216 mm, resolution = 2.4 x 2.4 x 2.4 mm. The full details of the imaging acquisition protocol are described in [24].

### C. Inter-modality source coupling

In this study, and motivated by prior work showing similarity between structural covariation and rs-fMRI networks [15, 25, 26], we adopted a standardized functional template as a spatial prior for our data to ensure consistency and comparability. For that, we applied a spatially constrained ICA independently to both rs-fMRI and sMRI. This approach allows the priors to be optimized and updated for individual subjects and modalities by maximizing spatial independence, while also allowing us to evaluate coupling among modalities. Using the resulting matrices from these two modalities, we evaluated their linkage via a similarity measure (cross-correlation) to assess the inter-modality source coupling. The IMSC analysis, focusing on the relationship between brain function and structure, allows us to evaluate the corresponding functional fingerprint of the whole brain structure.

For the ICA analysis, we first calculated a fully automated constrained ICA using the 53 component Neuromark_fMRI_1.0 template [27] independently on both the rs-fMRI and sMRI data to adapt the template to the data while also providing correspondence between networks. This step was performed using the GIFT toolbox [28]. For the derived rs-fMRI components, we then computed the functional network connectivity (FNC) [29, 30]. For the sMRI we calculated a source-based morphometry (SBM), estimating subject specific loading parameters corresponding to each of the 53 areas.

To evaluate the inter-modality source coupling we computed the cross correlation of the sMRI loading parameter vector with the FNC matrix for each subject. This analysis resulted in a matrix of 53 brain areas for each subject, labeled as IMSC matrix, that creates a link between a whole brain structural fingerprint and the corresponding functional fingerprint (see Fig. 1).

**Fig. 1.**
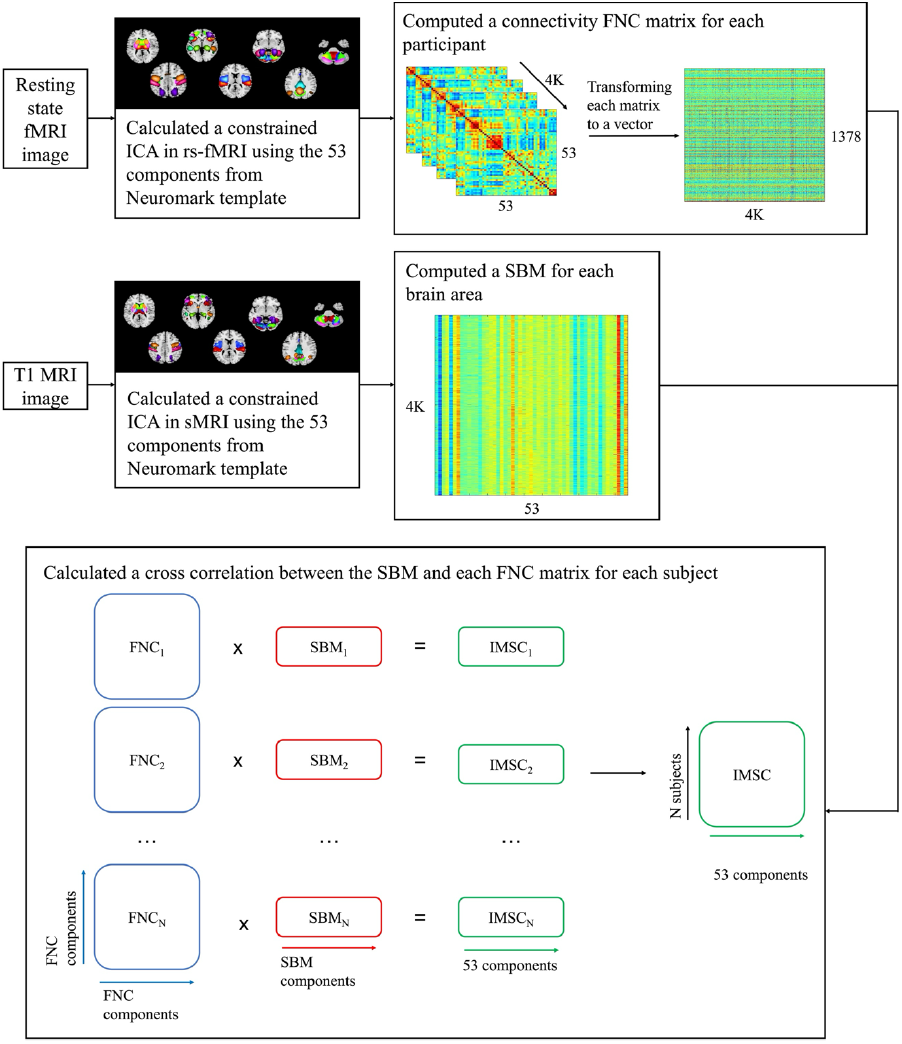
Inter-modality source coupling approach. This approach uses rs-fMRI and sMRI images and performs an independent constrained ICA on both images, using the same Neuromark spatial priors. For the rs-fMRI data, we computed a FNC matrix for each subject, this matrix represents the connectivity among the 53 brain regions, these matrices were then converted into a vector for each participant. For the sMRI it was calculated a SBM matrix that created a specific loading parameter vector for each subject (representing the covariation among subjects, or the degree to which each subject expresses the structural source). We then calculated a cross-correlation between the FNC and SBM matrices, resulting in the IMSC matrix. This matrix allows us to evaluate the relationship between brain function and structure.

### D. Evaluation of relationship to reading ability

Next, we computed the cross-correlation for the brain areas from the last analysis and a measure of reading ability (the individual ability score). This resulted in a vector of the areas showing an association between the structure-function fingerprint and reading ability. We obtained the overall average score for all four thousand participants (see Fig. 2). We also performed a replication in the independent subset with four thousand participants of the ABCD data. We present only the results that replicated.

**Fig. 2.**
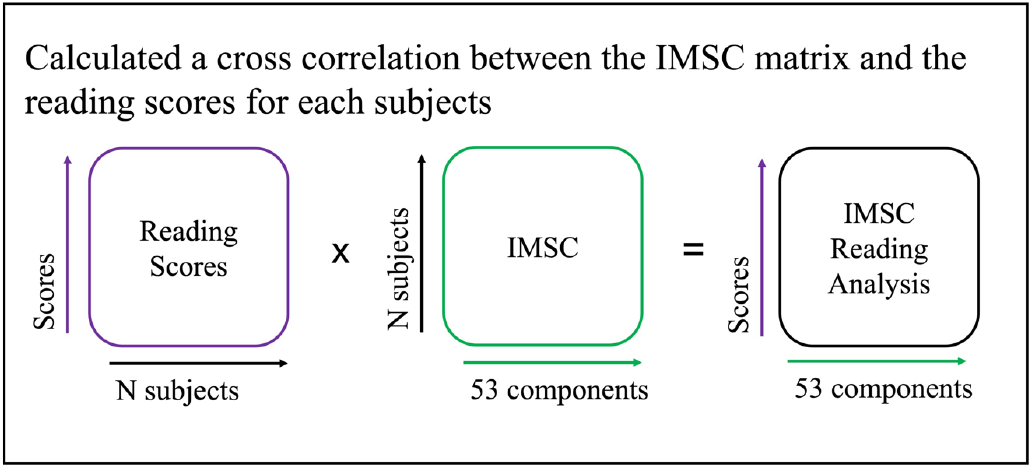
Inter-modality source coupling approach applied to study the relationship between IMSC and reading ability. We present the overall average analysis, where we calculated an IMSC matrix for each of the 53 components associated with the individual subjects that was then cross correlated with the respective reading score for each subject, resulting in an IMSC matrix linking each subject’s reading score to a distinct brain area.

## III. Results

This study unveiled a novel data-driven methodology for assessing the relationship between resting-state functional (rs-fMRI) and structural (sMRI) brain data. This study also evaluated how this relationship associated to the higher-level cognitive skill of reading, investigating how various linked multimodal brain regions underly proficient and struggling readers. Additionally, we generated group maps associated with rs-fMRI and sMRI data created from the constrained ICA, revealing a meaningful correspondence between them.

To facilitate a comparison between the two modalities for individual subjects, it was essential to generate independent, adaptative maps tailored to each subject for rs-fMRI and for sMRI. An ICA was performed on both rs-fMRI and sMRI, Fig. 3 displays the resulting group maps from these analyses. The groups maps are ICNs that were arranged in functional domains according to their functional and anatomical roles, the subcortical, auditory, sensorimotor, visual, cognitive control, default mode and cerebellar domains [31].

**Fig. 3.**
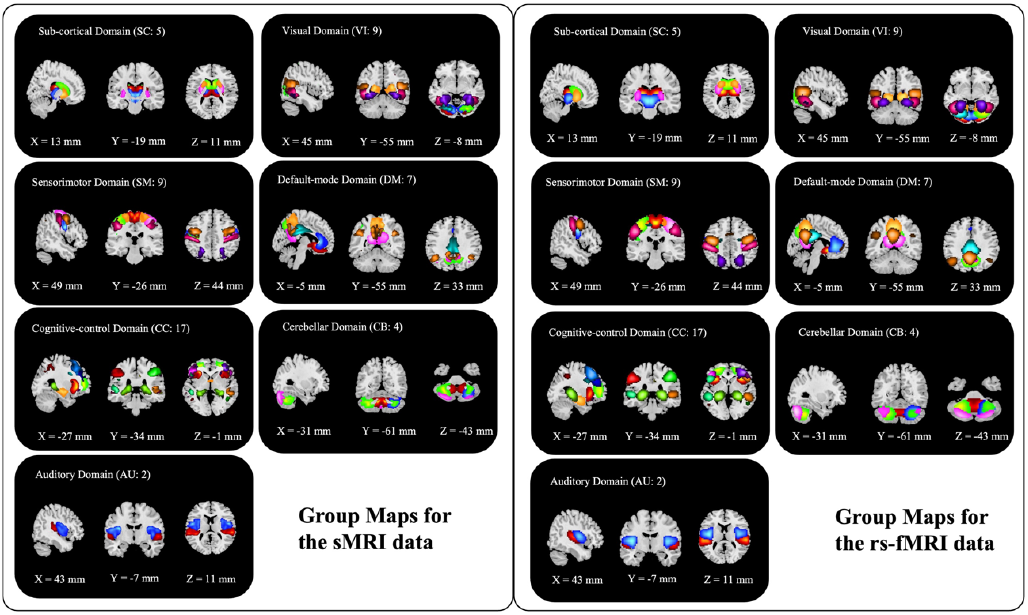
Group-level maps for the sMRI and rs-fMRI components. These show the group maps calculated using a constrained ICA and the same template with 53 different components for both rs-fMRI and sMRI data, where each color represents a specific component. Although differences can be observed between the domains, there is a strong correspondence between the two modalities due to the spatial constraint.

Generally, there is a good correspondence between the two modalities in the adaptive components. In the analysis of rs-fMRI data, we observe smoother patterns, whereas the sMRI data tend to exhibit more focal characteristics, showing slightly less lateralization. Notably, we identify some specific variations, including high activity in thalamus, in the subcortical domain, and precuneus, in the default mode domain, in rs-fMRI scan, in contrast to slightly lower covariation in sMRI images.

For the correlation analysis with linked brain areas using the results from the IMSC, we calculated p-values and selected 0.05 as significance cutoff after correcting for multiple comparisons. Results are presented below (see Table I) where we detail the structural/functional coupling brain areas associated with the individual ability score and indicate if the correlation observed was positive or negative. All the results presented below were replicated in a second analysis.

**TABLE 1.** Results for correlation between structural/functional coupling brain areas and the reading ability score.

| Correlation | Brain Areas |
| --- | --- |
| Positive | Inferior parietal lobule |
|  | Middle cingulum gyrus |
|  | Middle frontal gyrus |
|  | Superior frontal gyrus |
|  | Superior parietal lobule |
| Negative | Cuneus |
|  | Fusiform |
|  | Lingual |
|  | Middle temporal gyrus |

Results showing significant (and corrected for multiple comparisons) positive or negative correlations for specific structural/functional coupling brain areas associated to the individual ability reading score. It should be noted that higher level processing areas, such as the frontal and parietal lobe level of sMRI and rs-fMRI coupling are linked to increased reading scores while early processing areas, such as the cuneus and the lingual gyrus, sMRI and rs-fMRI coupling are linked to reduced reading scores.

In Fig. 4 we present the results from the analysis along the spatial maps highlighting significant brain areas. When assessing both positive and negative correlations, it became apparent that frontal brain areas exhibited important positive coupling with rs-fMRI/sMRI in relation to reading ability, while areas associated with early processing demonstrated a negative coupling to reading ability. (see Fig. 5).

**Fig. 4.**
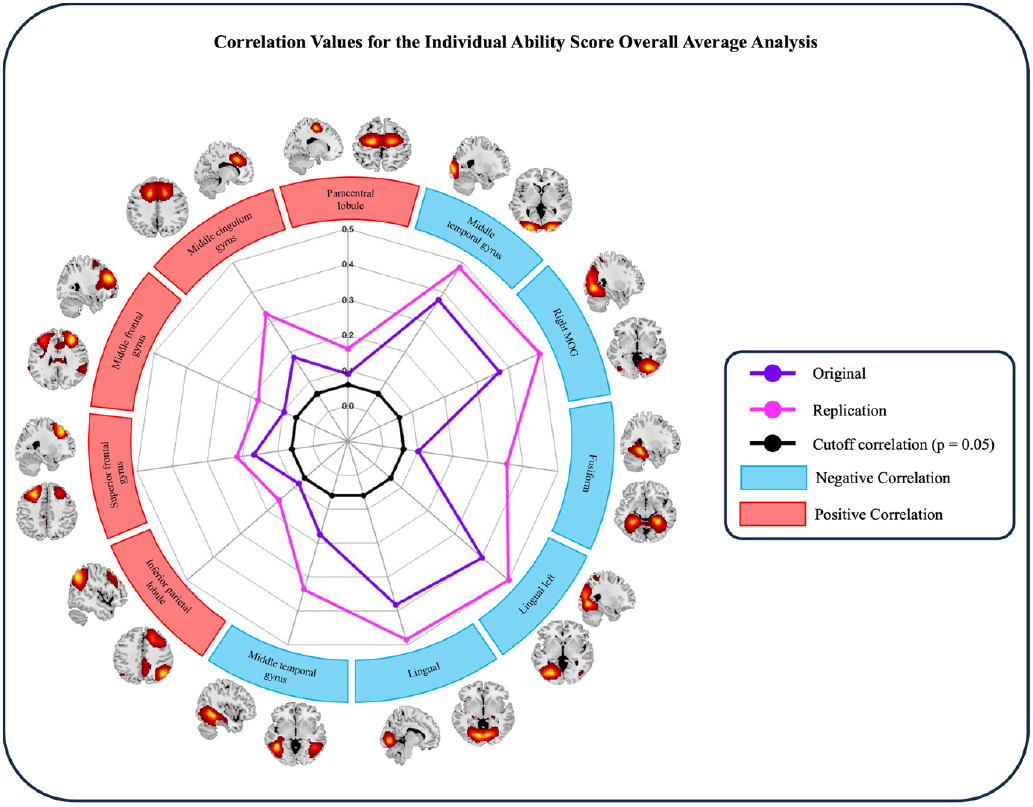
Plot showing the absolute correlation values for the regions that were significant for the analysis. The line in purple is for the original analysis and the pink line is for the replication. The blue are components that showed a negative correlation and the red are components that showed a positive correlation.

**Fig. 5.**
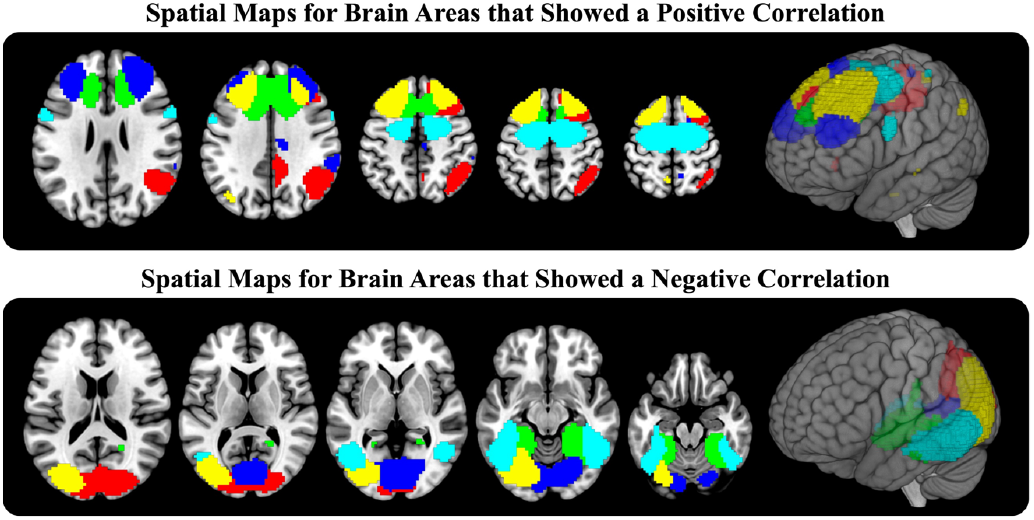
Horizontal brain slices and a rendered view for sMRI/rs-fMRI coupling showing components that exhibited significant positive (top) or negative (bottom) correlation with reading. It is noted that all positive correlations are focused in frontal, and superior and inferior parietal lobules, while negative correlation are focused in early processing areas, such as the fusiform and lingual gyrus.

## IV. Discussion

In this work we present a new multimodal analysis approach we are calling IMSC, which allows us to study the relationship of reading to the strength of coupling between brain structure and function in brain relevant networks. Our results suggest that the coupling of function and structure within multiple brain areas are associated with reading-related differences in children between the ages of 9 and 11. Many of these identified areas have previously been implicated using unimodal approaches in understanding the neural basis of reading ability and related language processing [32-35]. This connection to previous research highlights the importance and significance of our findings in understanding the broader context of cognitive neuroscience. These coupled brain areas could represent a potential key to understanding the neural basis of developing reading and language skills. The understanding of the specific development and function of these regions could pave the way for more effective strategies to enhance reading abilities and related language processing. Moreover, this newly proposed method demonstrated robust and replicable results in different subsets of subjects, suggesting valuable insights into these intricate brain interconnections. Our IMSC method also stands out due to its utilization of the same brain template priors for both rs-fMRI and sMRI, streamlining the comparison process and offering a standardized framework for more accurate and meaningful analyses.

Based in the modern vision of the cortical networks for reading [36], we observed higher reading abilities and an increased structure-function coupling in areas related to access to meaning, pronunciation and articulation. Also, with higher reading abilities we observed a decreased structure-function coupling in the visual word-form area and visual inputs, but increased coupling with speech related areas in the frontal brain areas [36]. We also observed a significant correlation between higher reading abilities in participants and an increased coupling between function and structure in the frontal brain areas, such as the inferior parietal lobule and superior frontal gyrus, both of which are widely recognized as a key component of the language and reading processing networks [32, 35]. Our observations also revealed that participants exhibiting lower reading abilities demonstrated a higher coupling between function and structure in the lingual gyrus, that are involved in the early processing of letters before significant linguistic processing [37]. Overall, our study unveils a plausible relationship between reading and multimodal brain structure/function. Specifically, we observed a positive relationship between reading and the multimodal linkage among higher order cognitive brain regions, such as the inferior parietal lobule and the superior frontal gyrus, shedding light on the higher-level cognitive processes intricately linked to proficient reading abilities. This positive multimodal association suggests an interesting link between the complex cognitive challenges and multiple brain networks’ involvement intrinsic to proficient reading. Conversely, we identified a negative relationship between reading and more basic sensory/perceptual regions, such as the middle occipital gyrus, suggesting that as reading proficiency increases, there is a corresponding decrease in the structure and functional coupling of sensory processing areas. This more complex understanding of the relationships between reading and distinct, multimodal brain regions begins to contribute to an increasingly sophisticated comprehension of the neural structure underlying reading skills, emphasizing the complex interconnections between cognitive and sensory domains, and the level of coupling between brain functions and structures.

It is worth noting that our study also revealed a good correspondence between the spatial networks from the constrained ICA derived from both rs-fMRI and sMRI data. Results show more granularity in the sMRI networks, due to the higher spatial resolution. In addition, there is a good correspondence suggesting an interplay between brain structure and function in our participants. It is worth noting that, though the ICA approach does introduce a strong spatial prior, we have previously validated the approach and find that it does a good job in reflecting the underlying data, that is, if there are not structural networks present, the results would be very unlikely to show such a result.

Whereas we sought to enhance the robustness of our findings by replicating our analyses using a different subset of the ABCD data, future studies should extend our replication by using data from different studies. Additionally, it is worth noting that though our use of rs-fMRI templates as a prior in conjunction with sMRI data is reasonable given our prior observations of resting network properties present in covarying sMRI data, there are alternatives that can be explored. For example, a promising direction for future work could involve exploring the use of sMRI templates and investigating the potential linkages between the two imaging modalities. Furthermore, an important area for exploration is the examination of changes in intrinsic coupling over time, such as during children’s reading development over multiple ages. Future studies may delve into the longitudinal aspects of our findings by exploring alterations in intrinsic coupling measured by IMSC over extended periods. Another worthwhile exploration could leverage the IMSC coupling method to examine different forms of brain coupling in other relevant contexts besides reading. An additional potential limitation is the reliance on a single reading ability test for assessing participants’ reading performance. Subsequent studies could enhance the robustness of the analysis by incorporating various tests to ensure a comprehensive evaluation of reading abilities. These considerations lay the groundwork for future investigations to expand and refine our understanding of the links between brain structure, brain function, and reading.

## V. Conclusion

In summary, we proposed a new method called inter-modality source coupling (IMSC) to study the level of sMRI and rsMRI coupling and associated it to reading ability in children. Our findings indicate that heightened sMRI and rsMRI networks coupling within higher cognitive brain regions correlates with elevated reading scores, while coupling within early-processing sensory or perceptual networks was linked to lower reading scores. This supports current thinking about a potential hierarchy in the neural mechanisms involved in reading, where higher cognitive regions, such as the inferior parietal lobule and middle frontal gyrus, play key roles that may contribute to proficient reading skills, while altered coupling patterns in early-processing networks, such as the lingual gyrus, may be indicative of challenges in reading acquisition or comprehension. Further investigation into those coupling patterns could provide valuable insights into the nuanced, but complex relationship between structure/function brain coupling and reading abilities.

The alignment of our results with the modern vision of cortical networks for reading underscores the relevance of our study in the broader context of cognitive neuroscience. Overall, we linked a pattern of whole brain structure for a specific child to the corresponding whole brain functional connectivity pattern for a specific network and our results revealed significant associations between the coupling of specific brain areas and reading performance. Future studies will need to further our understanding of how such coupling changes during development and how it may impact new learning and the development of reading and other related higher-order cognitive abilities. This study suggests that the use of IMSC has potential to further the development more sophisticated data-driven models of such complex neurocognitive systems.

